# SMRT sequencing generates the chromosome-scale reference genome of tropical fruit mango, *Mangifera indica*

**DOI:** 10.1101/2020.02.22.960880

**Authors:** Wei Li, Xun-Ge Zhu, Qun-Jie Zhang, Kui Li, Dan Zhang, Cong Shi, Li-Zhi Gao

**Affiliations:** Institution of Genomics and Bioinformatics, South China Agricultural University, Guangzhou 510642, China; Plant Germplasm and Genomics Center, Germplasm Bank of Wild Species in Southwestern China, Kunming Institute of Botany, Chinese Academy of Sciences, Kunming 650204, China; University of Chinese Academy of Sciences, Beijing 100039, China; School of Life Sciences, Nanjing University, Nanjing 210093, China; Novogene Bioinformatics Institute, 100083 Beijing, China

## Abstract

Mango (*Mangifera indica*), a member of the family Anacardiaceae, is one of the world’s most popular tropical fruits. Here we sequenced the variety, “Hong Xiang Ya”, and generated a 371.6-Mb mango genome assembly with 34,529 predicted protein-coding genes. Aided with the published genetic map, for the first time, we assembled the *M. indica* genome to the chromosomes, and finally about 98.77% of the genome assembly was anchored to 20 pseudo-chromosomes. The availability of the chromosome-length genome assembly of *M. indica* will provide novel insights into genome evolution, understand the genetic basis of specialized phytochemical composites relevant to fruit quality, and enhance allele mining in genomics-assisted breeding for mango genetic improvement.

## Background & Summary

Mangos, renowned as “king of fruits”, are native to South and Southeast Asia but are now enjoyed all over the world1. The worldwide production volume of mangos, mangosteens, and guavas reached 50.65 million metric tons in 2017, an increase from 46.5 million metric tons in the 2016, which globally grades as the fifth most produced fruit crop. Much of the world’s mangos are produced in the Asia Pacific region, such as India, China and Thailand (http://www.fao.org/faostat/). The genus *Mangifera*, which belong to the Anacardiaceae family, comprise about 50 species^2^. While other *Mangifera* species may also yield edible fruits of mangos, almost all cultivated mangoes come from only one species, *Mangifera indica*^3^. This species possesses a long domestication and cultivation history of over 4,000 years in the Indo-Burmese and Southeast Asia regions, and then spread to other areas of the world since the 14^th^ century^4,5^. This famous fruit tree is extensively grown in tropical regions and extend to subtropical regions in the world nowadays.

While some mango fruits are often processed into various kinds of products, for example, nectar, juice and jam, the majority are predominantly consumed in fresh^1^. As many other Anacardiaceae plants, mangos contain phenolic compounds that may stimulate interaction dermatitis, an undesired characteristic for numerous users of fresh mango fruits^6^. However, this fruit with gorgeous appearance and unusual flavors has attracted an increasingly large number of world consumers. Nevertheless, the biosynthesis of these compounds remain largely unresolved up to date. Furthermore, although mango cultivars have heavily relied on vegetative propagation, recent decades have witnessed huge efforts in US, China, Australia and other countries, through conventional cross-breeding programs. The generation of a large number of mango varieties have progressively speeded its wide-reaching distribution in the world^7^. Although some cytogenetics data^8^, genetic mapping^9,10^ and transcriptomics data^11,12^ are publicly available, the lack of chromosome-level high-quality mango reference genome sequences has seriously hampered our understanding of the genetic basis of specialized phytochemical composites and allele mining in genomics-assisted breeding for mango improvement.

The completion of chromosome-scale genome assembly of *M. indica* is able to build the foundation for the discovery and application of desired agronomic traits that is of great interest in modern genetic improvement program. Here, we report the first *de novo* assembled chromosome-level genome of *M. indica* using SMRT sequencing technology with the assistance of the published genetic map. The obtained genome assembly will greatly help to obtain novel insights into the genome evolution and metabolic biosynthesis of important compounds relevant to fruit quality and enhance the genomics-based trait improvement and mango germplasm utilization.

## Methods

### Sample collection, library construction and sequencing

An individual plant of the *M. indica* variety, “Hong Xiang Ya”, grown in Xishuangbanna City, Yunnan Province, China, was collected for the genome sequencing. Fresh and healthy leaves were harvested and immediately frozen in liquid nitrogen, followed by storage at −80°C in the laboratory prior to DNA extraction. High-quality genomic DNA was isolated using a modified CTAB method^13^ for both Illumina and Pacbio sequencing. The quantity and quality of the DNA sample were examined using a NanoDrop 2000 spectrophotometer (NanoDrop Technologies, Wilmington, DE) and electrophoresis on a 0.8% agarose gel, respectively. For Illumina sequencing, the extracted genomic DNA was fragmented using S220 Focused-ultrasonicator system (Covaris Inc, USA). One paired end library was constructed following the Illumina’s instructions and sequenced on Illumina Hiseq4000 platform. For Pacbio sequencing, a 40-kb SMRTbell DNA library was prepared and sequenced on PacBio Sequel II platform with one SMRT cell. Finally, a total of 87.67 Gb short sequencing data and 151.09 Gb Pacbio data were generated, respectively (Table 1).

**Table 1.**
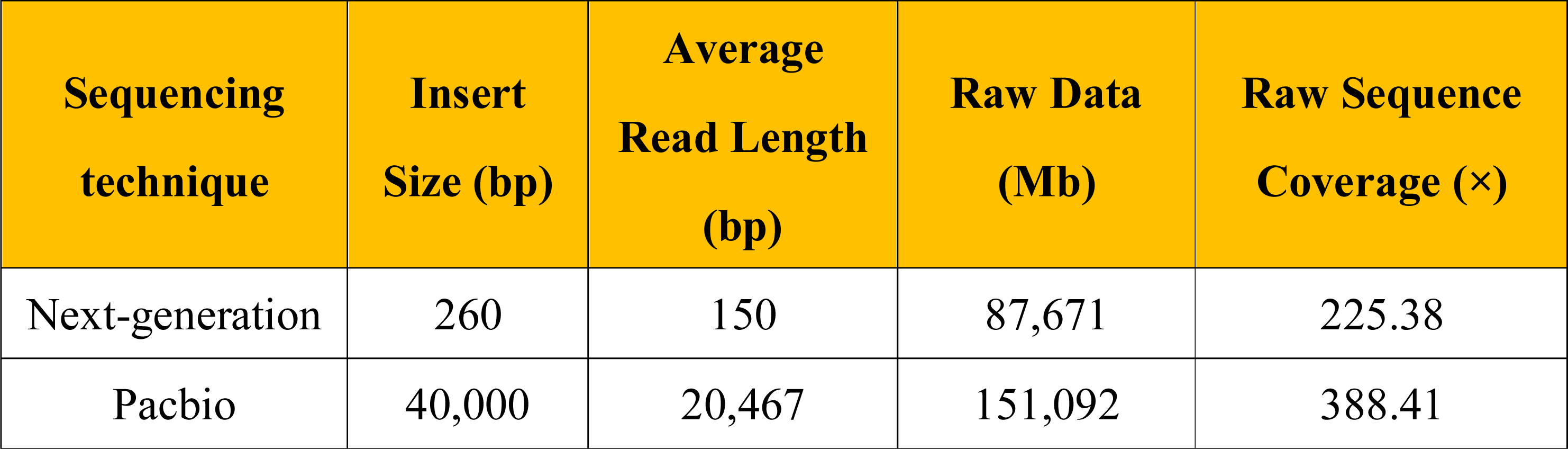
Libraries and read statistics used for the mango genome assembly. The estimated genome size is ∼389 Mb.

### Estimation of genome size and heterozygosity

The genome size of mango was estimated using *k*-mer frequency distribution generated from short reads. Jellyfish^14^ was used to calculate 17-mer abundance, and genome size of mango was then estimated using the formula: Genome size = *k*-mer_num/peak_depth, where *k*-mer_number is the total number of *k*-mer and peak_depth is the peak value of *k*-mer frequency distribution. In this study, a total of 43,154,105,487 *k*-mer were counted, while the main peak occurred at depth 111 corresponds to unique haploid sequences. The genome size of mango was estimated to be ∼389 Mb. A small peak was also detected at 1/2 peak depth corresponding to considerable heterozygous fractions (Fig. 1). Genomescope^15^ was then used to estimate the heterozygosity level of mango. The *k*-mer histogram generated by Jellyfish^14^ was subjected to Genomescope^15^ (http://qb.cshl.edu/genomescope/) and heterozygosity estimate for the mango genome was in the range of ∼1.84-1.85%.

**Figure 1.**
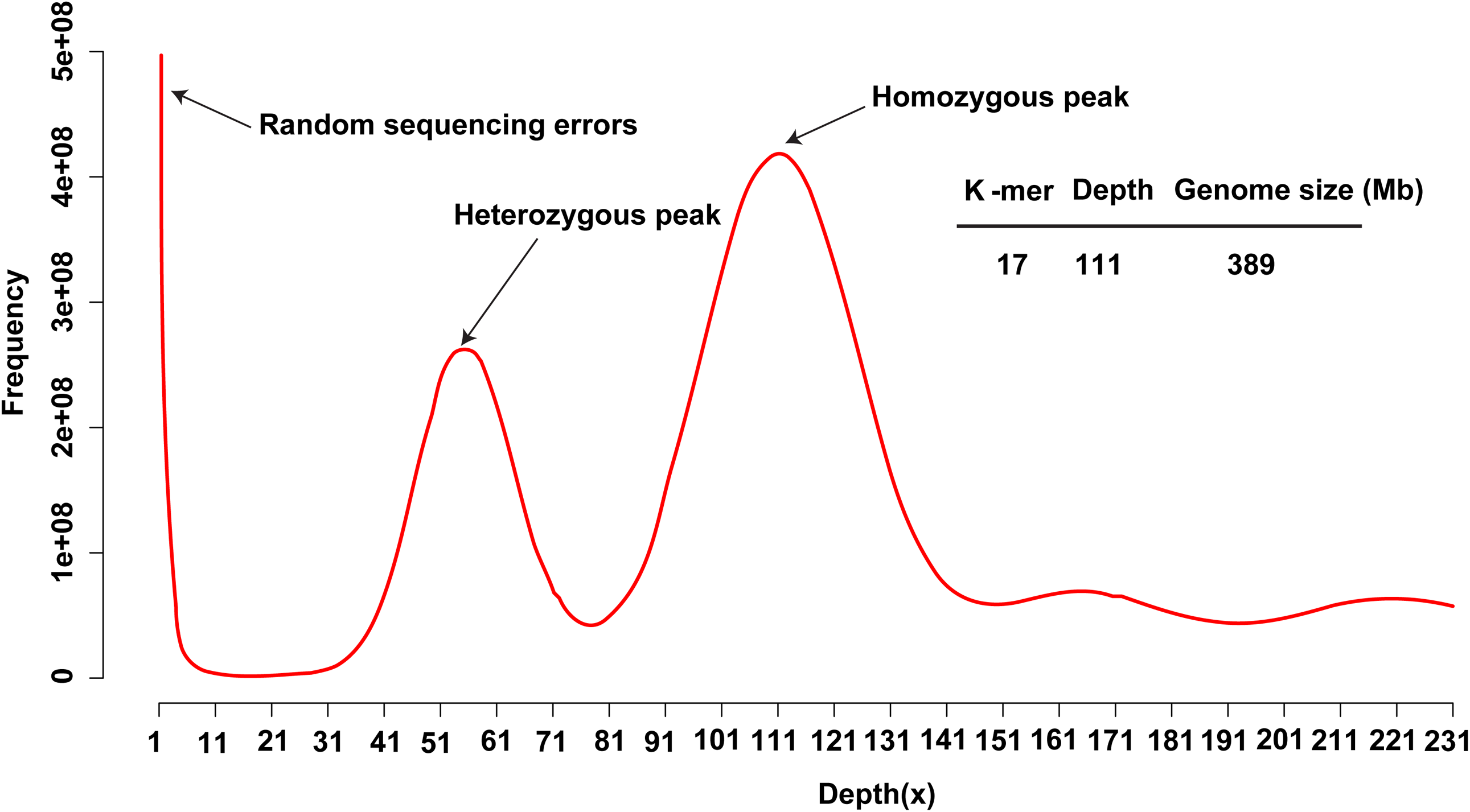
The 17-*mer* distribution of sequencing reads of the mango genome. The occurrence of 17-*mer* was calculated using jellyfish based on the sequencing data from short insert size libraries (insert size ≤ 500 bp) of mango. The sharp peak on left with low depths represents the essentially random sequencing errors. The middle and right peaks indicate the heterozygous and homozygous peaks, the depths of which are 55 and 111, respectively.

### *De novo* genome assembly

In an attempt to minimize the effect of high heterozygosity, Falcon and Falcon-unzip^16^ were used to perform the genome assembly. The longest 55× subreads were selected as seed reads to perform interactive error correction, Falcon was used to obtain primary contigs (p-contigs). The p-contigs were then phased using Falcon-Unzip. Two subsets of contigs were generated, including the primary contigs (p-contigs) and the haplotigs, which represent divergent haplotypes in the assembly. Both p-contigs and haplotigs were polished as follows: firstly, all the pacbio reads were aligned against the assembly using pbalign (https://github.com/PacificBiosciences/pbalign). The output files were fed to Arrow (https://github.com/PacificBiosciences/GenomicConsensus) implemented in SMRTLink to polish the assembly. Next, the Illumina data from short libraries were aligned to the polished assembly using BWA^17^ with default parameters, and then, Pilon^18^ was used for sequence assembly refinement based upon these alignments. Two rounds of Pilon was performed to correct the single-base errors and small indels in the assembly. To remove the allelic contigs retained in primary assembly which is deleterious for downstream analysis, Purge Haplotigs^19^ was used to identify and reassign allelic contigs. Briefly, the pacbio raw reads were mapped to the p-contigs using Minimap2^20^. The resulting BAM file was used to generate a read-depth histogram. The collapsed haplotype contigs will fall into the 1×read-depth peak, whereas the allelic contigs will result in half the read-depth. Finally, we generated a primary assembly with a total length of ∼372 Mb which spanned 95.37% of the genome size estimated by *k*-mer analysis (Table 2). The assembly comprised of 120 contigs with an N50 size of 4.82 Mb (Table 2). We also generated a combined 168.20 Mb of haplotype-resolved sequence, with an N50 of 458.69 Kb and a maximum length of 2,007.74 Kb (Table 2). The published genetic map^9^ was also used to anchor scaffolds to chromosomes with ALLMAPS^21^. A total of 112 contigs were anchored to 20 pseudochromosomes (Table 2). The anchored contigs were 367.05 Mb in size, occupying 98.77% of the genome (Table 2). The chromosome lengths of the mango genome varied from ∼12 Mbp (Chr10) to ∼28 Mbp (Chr19) with an N50 size of ∼19 Mbp (Table 3; Fig. 2).

**Table 2.**
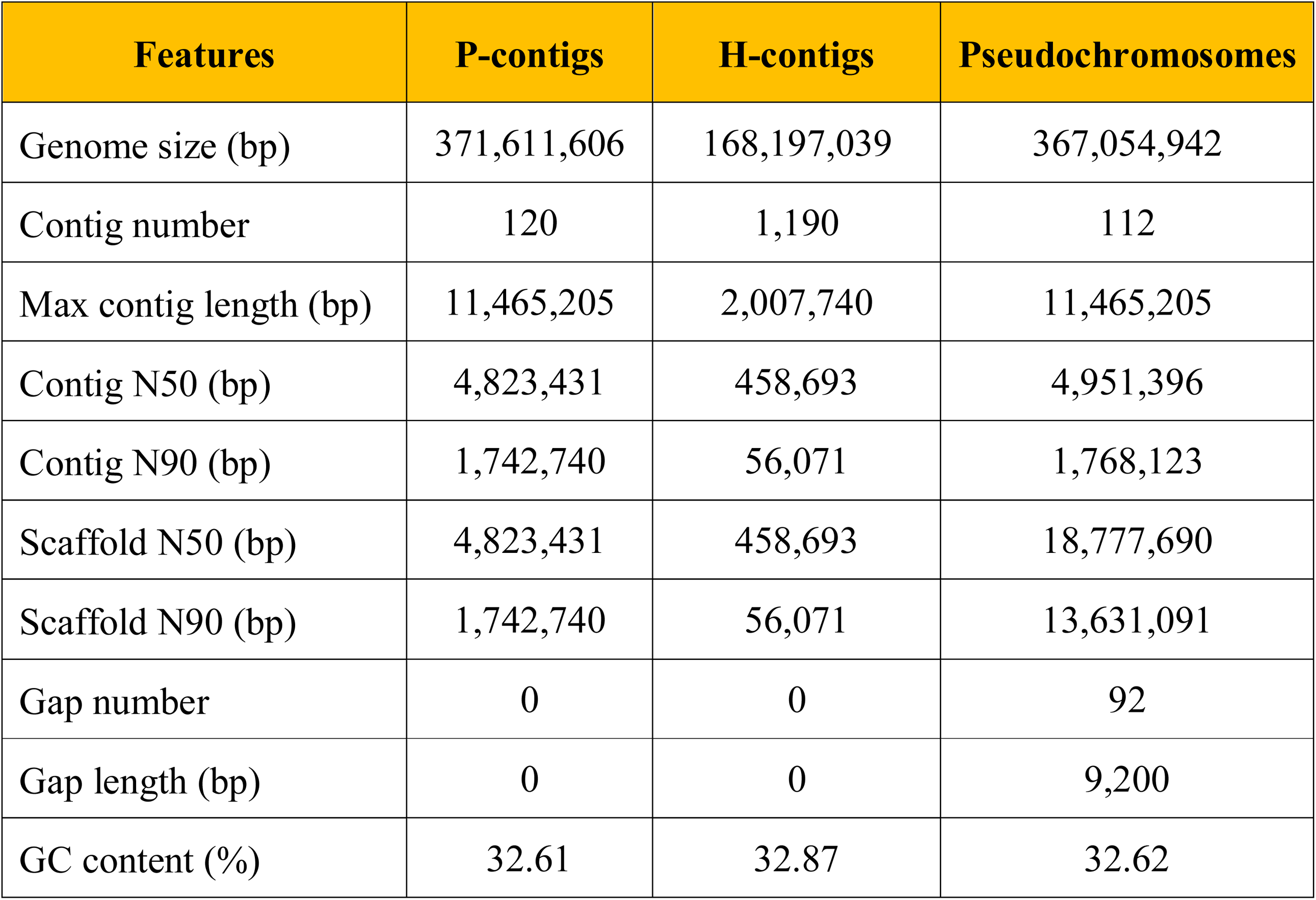
Assembly statistics of the mango genome.

**Table 3.**
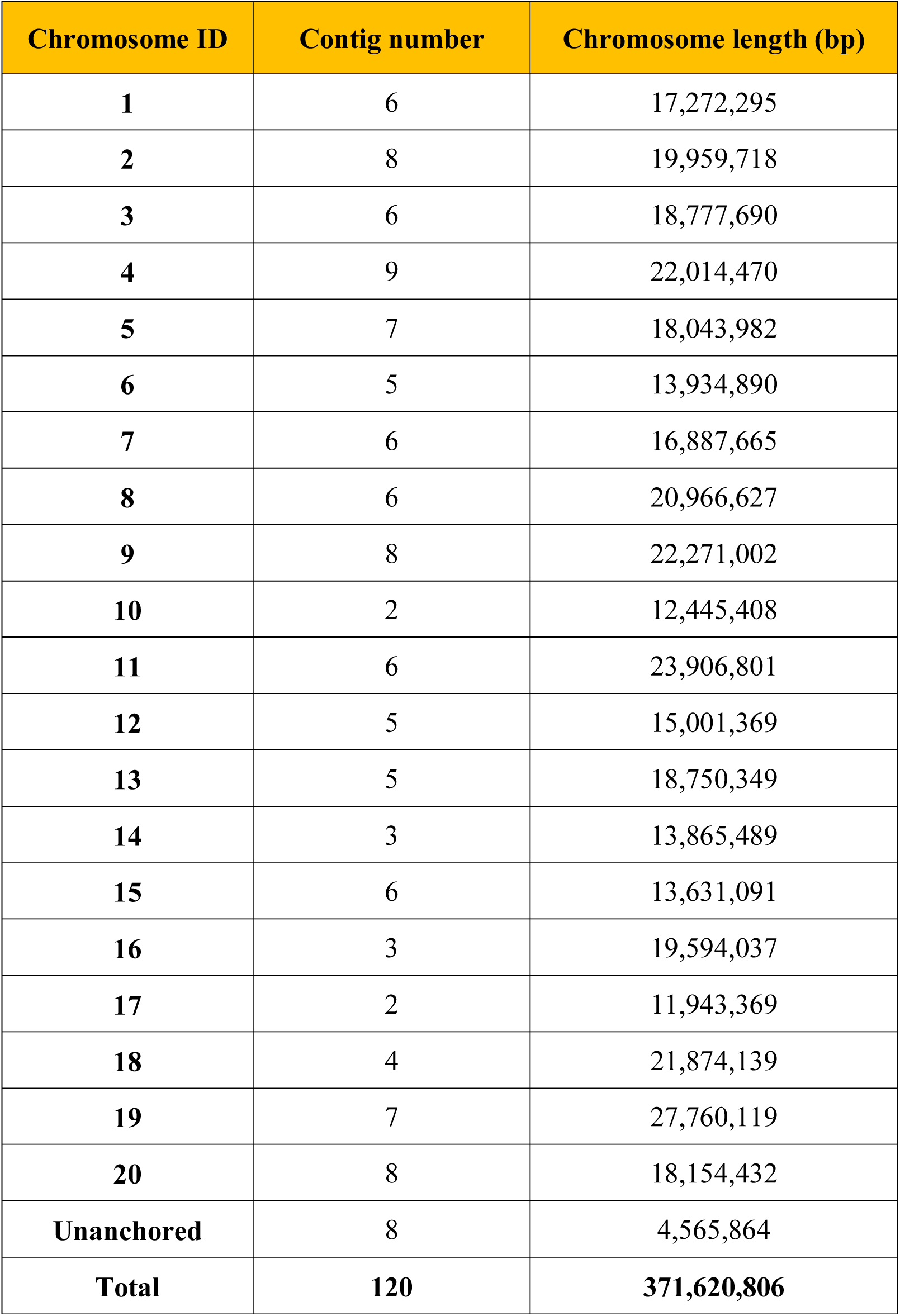
Chromosome lengths of the assembled mango genome.

| Chromosome ID | Contig number | Chromosome length (bp) |
| --- | --- | --- |
| **1** | 6 | 17,272,295 |
| **2** | 8 | 19,959,718 |
| **3** | 6 | 18,777,690 |
| **4** | 9 | 22,014,470 |
| **5** | 7 | 18,043,982 |
| **6** | 5 | 13,934,890 |
| **7** | 6 | 16,887,665 |
| **8** | 6 | 20,966,627 |
| **9** | 8 | 22,271,002 |
| **10** | 2 | 12,445,408 |
| **11** | 6 | 23,906,801 |
| **12** | 5 | 15,001,369 |
| **13** | 5 | 18,750,349 |
| **14** | 3 | 13,865,489 |
| **15** | 6 | 13,631,091 |
| **16** | 3 | 19,594,037 |
| **17** | 2 | 11,943,369 |
| **18** | 4 | 21,874,139 |
| **19** | 7 | 27,760,119 |
| **20** | 8 | 18,154,432 |
| **Unanchored** | 8 | 4,565,864 |
| **Total** | **120** | **371,620,806** |

**Figure 2.**
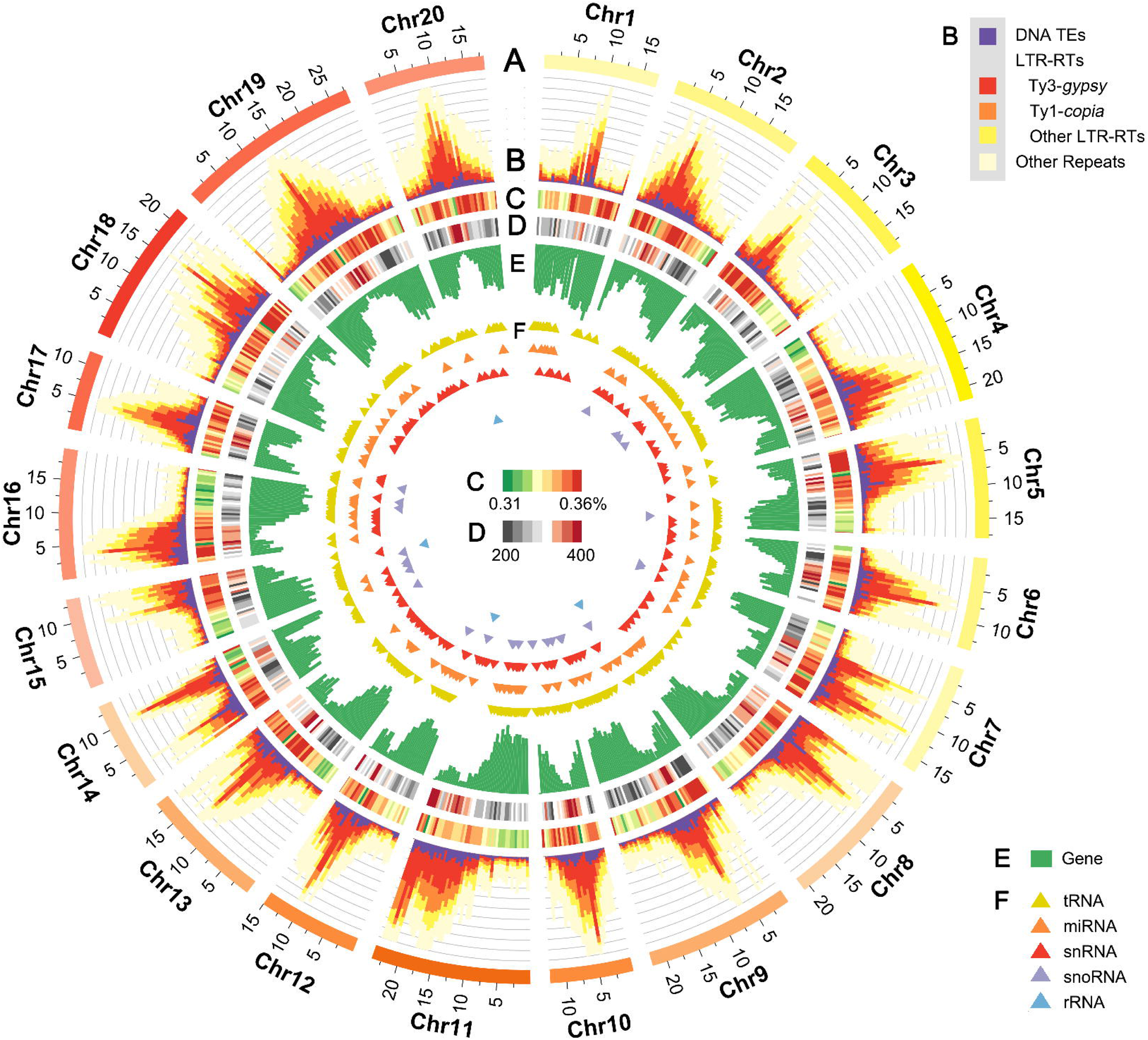
The mango genome features. (A) Circular representation of the 20 pseudochromosomes; (B) the distribution of TEs. Purple for DNA TEs, red for Ty3-*gypsy* LTR-RTs, orange for Ty1-*copia* LTR-RTs, yellow for unclassified LTR-RTs. and light yellow for other repeats; (C) GC content; (D) number of SSRs per 500kb; (E) the distribution of protein-coding genes; (F) the distribution of non-coding RNAs, yellow for tRNAs, orange for miRNAs, red for snRNAs, purple for snoRNAs, and blue for rRNAs.

### Transposable element annotation

A mango-specific repeat library was constructed following the instructions of “Repeat Library Construction-Advanced” (http://weatherby.genetics.utah.edu/MAKER/wiki/index.php/Repeat_Library_Construction-Advanced) in MAKER-P pipeline^22^. Firstly, the genome assembly was searched for miniature inverted transposable elements (MITEs) using MITE-Hunter^23^. The identified MITEs were manually checked for TSD and TIR. Long terminal repeat retrotransposons (LTR-RTs) were identified using LTRharvest^24^ (implemented in GenomeTools^25^) and LTR_Finde^r26^. LTR_retriever pipeline^27^ was applied to integrate the results of LTR_Finder^26^ and LTRharvest^24^ to efficiently remove false positive elements. Repeatmodeler (http://www.repeatmasker.org/RepeatModeler/), which can automatically execute two core *de novo* repeat finding programs, including RECON2^8^ and RepeatScout^29^, was also used to construct repeat library. The elements identified as unknown were searched against the transposase database. An in-house perl script was used to include the sequences matching transposase into the relevant superfamily. All the identified repeat libraries were searched against a plant protein database where proteins from transposons were excluded. Elements with significant hits were filtered using ProtExcluder^22^. The remained elements were merged to form a comprehensive TE library. Repeatmasker^30^ was used for the annotation of repetitive sequence. We also identified the tandem repeats using the Tandem Repeat Finder (TRF) package31 and the non-interspersed repeat sequences using RepeatMasker^30^ with NCBI search engine. Results showed that approximately 47.94% of the mango genome consists of transposable elements (TEs). LTR retrotransposons were the most abundant TE type, occupying roughly 23.08% of the mango genome (Table 4). Most LTRs were LTR/*Gypsy* elements, which occupied 9.16% of the genome (Table 4). In addition, 18.12% of the mango genome comprised unclassified repetitive sequences (Table 4).

**Table 4.** Statistics of repeat sequences in the mango genome.

| Transposable Elements | Length (bp) | Percentage (%) |
| --- | --- | --- |
| **DNA transposons elements** | 16,187,808 | 4.36 |
| CMC | 157,712 | 0.04 |
| Harbinger | 89,828 | 0.02 |
| hAT | 3,270,678 | 0.88 |
| MuLE | 10,873,608 | 2.93 |
| Helitron | 1,713,184 | 0.46 |
| Others | 82,798 | 0.02 |
| **RNA transposons elements** | 88,061,304 | 23.70 |
| Non-LTR Retrotransposons | 2,297,579 | 0.62 |
| LINEs | 2,297,579 | 0.62 |
| LTR Retrotransposons | 85,763,725 | 23.08 |
| *Copia* | 24,945,038 | 6.71 |
| *Gypsy* | 34,039,343 | 9.16 |
| Caulimovirus | 418,105 | 0.11 |
| Others | 26,361,239 | 7.09 |
| **Unclassified elements** | 67,335,389 | 18.12 |
| **Other Repeats** | 6,580,619 | 1.77 |
| Low complexity | 906,641 | 0.24 |
| Simple repeats | 5,673,978 | 1.53 |
| **Total** | **178,165,120** | **47.94** |

Simple sequence repeats (SSRs) were identified in the mango genome using the MISA^32^ perl script with the settings: monomer (one nucleotide, n ≥ 12), dimer (two nucleotides, n ≥ 6), trimer (three nucleotides, n ≥ 4), tetramer (four nucleotides, n ≥ 3), pentamer (five nucleotides, n ≥ 3), and hexamer (six nucleotidess, n ≥ 3). In total, 284,679 SSRs were found, which constitutes 1.1% (4.07 Mb) of the genome (Table 5). Among the SSRs, monomer were most dominant (38% of the total SSRs), followed by dimer (26%), tetramer (16%), trimer (11%), pentamer (7%), and hexamer (3%), respectively (Table 5).

**Table 5.** Occurrence of simple sequence repeats (SSRs) in the mango genome.

| **Repeat type** | **Total counts** | **Total length (bp)** | **Average length (bp)** | **Proportion (%)** | **Frequency (loci/Mb)** | **Density (bp/Mb)** |
| --- | --- | --- | --- | --- | --- | --- |
| **Monomer** | 106,814 | 1,324,805 | 12 | 37.52 | 287.13 | 3,561.30 |
| **Dimer** | 72,740 | 1,220,894 | 17 | 25.55 | 195.54 | 3,281.97 |
| **Trimer** | 31,735 | 447,825 | 14 | 11.15 | 85.31 | 1,203.83 |
| **Tetramer** | 45,874 | 608,212 | 13 | 16.11 | 123.32 | 1,634.98 |
| **Pentamer** | 20,350 | 326,085 | 16 | 7.15 | 54.70 | 876.57 |
| **Hexamer** | 7,166 | 138,234 | 19 | 2.52 | 19.26 | 371.60 |
| **Total** | **284,679** | **4,066,055** | **14** | **100** | **765.27** | **10,930.26** |

### Gene prediction and functional annotation

Three independent approaches, including the *de novo* method, the homology-based method and the EST-aided method, were used to perform gene prediction. Augustus^33^ and SNAP^34^ were used to perform the *de novo* prediction, with two rounds of iterative training. The protein sequences from *Arabidopsis thaliana* (TAIR10)^35^, *Anacardium occidentalie* (phytozome v12), *Acer yangbiense*^36^ and *Citrus sinensis*^37^, were mapped to the genome by Exonerate^38^, using the Protein2Genome model. For RNA-seq aided gene annotation, the transcriptomic reads of five tissues, including seeds, mesocarps, leaves, flowers, and exocarps, were downloaded from NCBI SRA database under accession number SRR2163402- SRR2163406. Paired-end raw reads were trimmed using Trimmomatic^39^ to remove adaptors, reads with >3% N and low-quality reads. The quality-filtered reads were subjected to Trinity^40^ to perform *de novo* transcriptome assembly. The resulting transcripts were then aligned to the soft-masked mango genome using GMAP^41^ and BLAT^42^. The potential gene structures were iteratively refined using PASA (Program to Assemble Spliced Alignments)^43^. Finally, EVidenceModeler^44^ was used to integrate the predictions and generate a consensus gene set. Weights of evidences were manually set as: *ab initio* predictions, Augustus = 1, SNAP = 1; protein alignments, Exonerate = 4; EST-aided, PASA = 8. The gene set was refined with TransposonPSI^44^ to remove the TE-related gene model. Gene models with premature termination and/or consisting of fewer than 50 amino acids were also discarded. A total of 34,529 protein-coding genes with an average length of 3,295 bp were identified in the mango genome (Table 6). The average CDS length and exon number per gene were 1,143 and 5.6, respectively (Table 6).

**Table 6.** Prediction of protein-coding genes in the mango genome.

| Gene set |  | Number | Average length (bp) | Average CDS length (bp) | Average intron length (bp) | Average exon per gene |
| --- | --- | --- | --- | --- | --- | --- |
| *De novo* | Augustus | 33,485 | 2,709 | 1,203 | 330 | 5.6 |
|  | SNAP | 48,490 | 4,316 | 801 | 792 | 5.4 |
| Homolog | *Arabidopsis thaliana* | 15,268 | 3,138 | 1,099 | 428 | 5.8 |
|  | *Anacardium occidentale* | 117,919 | 3,860 | 1,247 | 479 | 6.5 |
|  | *Acer yangbiense* | 13,434 | 3,321 | 1,264 | 477 | 5.3 |
|  | *Citrus sinensis* | 26,222 | 3,791 | 1,320 | 459 | 6.4 |
| RNA-seq | PASA | 100,052 | 1,988 | 1,079 | 451 | 3.0 |
| Final set | EVM | 34,529 | 3,295 | 1,143 | 465 | 5.6 |

The predicted genes were searched against Swiss-Prot^45^ database using BLASTP (e-value cutoff of 10−5). The motifs and domains within gene models were identified using InterProScan^46^. Gene Ontology terms for each gene were directly retrieved from the corresponding InterPro entry. KAAS^47^ web server was applied to perform pathway analysis. All protein sequences were scanned with PfamScan^48^ with Pfam-A database^49^. A total of 90.71% gene models showed significant similarities to sequences in the public databases (Table 7).

**Table 7.** Functional annotation of the mango genome.

| Database | Number | Percentage (%) |
| --- | --- | --- |
| Total | 34,529 | 100 |
| InterPro | 31,143 | 90.19 |
| GO | 20,298 | 58.79 |
| KEGG | 8,731 | 25.29 |
| Pfam | 24,846 | 71.96 |
| Swissprot | 25,834 | 74.82 |
| Unannotated | 3,208 | 9.29 |

### Non-coding RNA gene annotation

The five different types of non-coding RNA genes, including transfer RNA genes (tRNA), ribosomal RNA genes (rRNA), small nucleolar RNA genes (snoRNAs), small nuclear RNA genes (snRNAs) and microRNA genes (miRNAs), were predicted in the mango genome. The tRNA genes were identified using tRNAscan-SE^50^. RNAmmer^51^ was used to predict rRNA genes and their subunits. For the identification of snoRNA, snoScan^52^ was used with the yeast rRNA methylation sites and yeast rRNA sequences provided by the snoScan distribution. Both snRNA genes and miRNA genes were identified by INFERNAL^53^ software against the Rfam database^54^. In total, we identified 598 tRNA genes, 45 rRNA genes, 47 snoRNA genes, 200 snRNA genes, and 235 miRNA genes, respectively (Table 8).

**Table 8.** Statistics for non-coding RNA genes in the mango genome.

| Non-coding RNAs | Number | Average Length (bp) | Total Length (bp) |
| --- | --- | --- | --- |
| tRNA | 598 | 74.13 | 44,332 |
| rRNA (8S) | 41 | 113.29 | 4,645 |
| rRNA (18S) | 1 | 1,808 | 1,808 |
| rRNA (28S) | 3 | 5,321 | 15,963 |
| snoRNA | 47 | 94.87 | 4,459 |
| snRNA | 200 | 127.04 | 25,408 |
| miRNA | 235 | 121.57 | 28,570 |

## Data Records

All the raw sequencing reads have been deposited to BIG Genome Sequence Archive database under accession number PRJCA002248. The genome assembly and genome annotation are also available at BIG Genome Warehouse under accession number PRJCA002248.

## Technical Validation

### Assessment of the genome assembly

First, high-quality reads from NGS sequencing were mapped to the genome assembly using BWA^17^. Our results revealed that nearly 88.19% Illumina reads were mapped to the genome assembly, among which 84.08% were properly mapped (Table 9). Second, the genome assembly was checked with benchmarking universal single-copy orthologs (BUSCO)^55^ from the Embryophyta lineage. Results showed a total of 1343 core orthologs (93.3%) could be found in the mango genome. Third, the RNA sequencing reads were assembled using Trinity^40^ and then aligned back to the assembled genome using GMAP^41^. Approximately 84.71% of the transcripts could be mapped to the genome (Table 9). Finally, we assessed the mango genome using the LTR Assembly Index (LAI)^56^, which evaluate the quality of assembly by the amount of identifiable intact LTR retroelements. The LAI score of mango is 13.05 (Fig. 3), indicating a high quality of genome assembly (draft quality, with LAI score less than 10; reference quality, with LAI score ranges from 10 to 20; and gold quality, with LAI score greater than 20).

**Table 9.** Quality assessment of the mango genome assembly and genome annotation.

|  | Methods | Total | Aligned | Percentage (%) |
| --- | --- | --- | --- | --- |
| **Genome Assembly** | **Reads mapping** |  |  |  |
|  | PE reads | 584,474,718 | 515,432,238 | 88.19 |
|  | **Transcripts of mango** |  |  |  |
|  | EST | 174,130 | 147,503 | 84.71 |
|  | **BUSCOs from Embryophyta lineage** |  |  |  |
|  | Complete | 1,440 | 1,343 | 93.26 |
|  | Duplicated | 1,440 | 237 | 16.46 |
|  | Fragmented | 1,440 | 29 | 2.01 |
|  | Missing | 1,440 | 68 | 4.72 |
| **Genome Annotation** | **Proteins from NCBI** | 532 | 469 | 88.16 |

**Figure 3.**
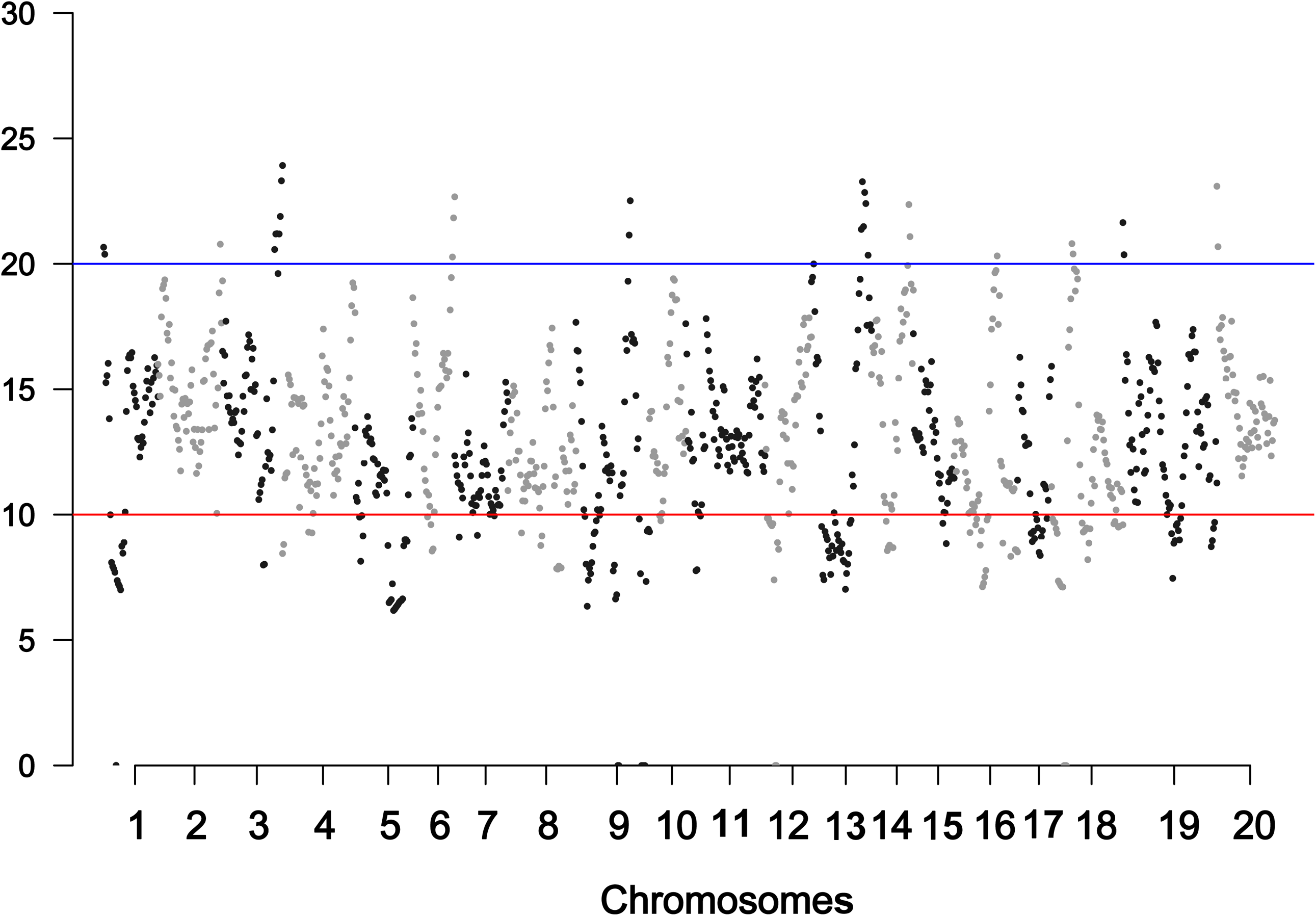
LAI scores in genomic regions of mango. Each dot represents LAI score with 300-Kb sliding window. The red line indicates the Draft-Reference boundary and the blue line shows the Reference-Gold boundary (Draft, such as Apple v1.0 and Cacao v1.0; Reference, such as Arabidopsis TAIR10; Gold, such as Rice MSUv7).

### Improvement of gene annotation quality

To improve the quality of gene prediction, we performed self-training with Augustus and SNAP. RNA-seq reads were *de novo* assembled using Trinity and refined with PASA to produce additional genome-guided transcriptome assemblies. Manual curation was performed with the training set, genes were retained if: (1) they have the complete gene structure without inner stop codons; they have multiple exons and the CDS length exceed 800 bp. CD-Hit^57^ was used to remove the training set with over 70% sequence similarity. Finally, a total of 2,738 high-quality gene models were filtered which served as the training sets. For Augustus, the traning sets were randomly divided into two parts, one with 2,238 gene models for training Augustus and the other with 500 gene models for assessing the accuracy of gene model. The protein sequences of mango were downloaded from National Center for Biotechnology Information (NCBI) database and plasmid-related proteins were removed. The protein sequences were then aligned against our gene predictions using BLAST program with an E-value cutoff of 1 × 10−10. Only hits with coverage >= 80% and identity >= 30% were retained. Results showed that 88.16% of the proteins were supported by our gene predictions, indicating that the annotated mango genes were of high quality (Table 9).

## Code availability

All the bioinformatics tools/packages used in this research described below along with their versions, settings and parameters.

**(1) Jellyfish:** version 2.2.10, -m 17 -s 200000M -t 20; **(2) GenomeScope**: k-mer_length=17, read_length=150; (**3) Falcon:** version 0.3.0, genome_size = 380000000, seed_coverage = 30, length_cutoff_pr = 5000, max_diff = 100, max_cov = 100; **(4) Falcon-Unzip:** version 0.3.0, default parameters; **(5) Pbalign:** version 0.3.1, --nproc 20; **(6) Arrow:** version 2.2.2, -j 50 --maskRadius 3; **(7) BWA:** version 0.7.12-r1039, default parameters; (**8) Pilon:** version 1.23, --fix snps, indels; **(9) Minimap2**: version 2.17-r974-dirty, -t 20 -ax map-pb --secondary=no; **(10) Purge Haplotigs: -**l 40 -m 130 -h 200; **(11) ALLMAPS:** default parameters; **(12) MITE-Hunter:** -n 5 -P 1 -S 12345678 -c 20; **(13) GenomeTools:** version 1.5.10, gt ltrharvest -similar 85 -vic 10 -seed 20 -seqids yes -minlenltr 100 -maxlenltr 7000-mintsd 4 -maxtsd 6 -motif TGCA -motifmis 1; (**14) LTR_Finder:** version 1.07, -D 15000 -d 1000 -L 7000 -l 100 -p 20 -C -M 0.9; **(15) LTR_retriever:** default parameters; **(16) RECON:** default parameters; (**17) RepeatScout**: version 1.05, default parameters; **(18) RepeatMasker:** version 1.332, -e ncbi; **(19) TRF**: version 4.09, default parameters; **(20) RepeatModeler:** version open-1.0.11, -e ncbi; **(21) MISA:** 1-12 2-6 3-4 4-3 5-3 6-3; (**22) Augustus:** version 2.7, --gff3=on; **(23) SNAP:** version 2006-07-28, default parameters; **(24) Exonerate:** version 2.2.0, --model protein2genome --minintron 20 --maxintron 30000 --showtargetgff --showvulgar 0 --showalignment 0 --softmasktarget TRUE --score 100 --percent 70; **(25) Trimmomatic**: version 0.32, LEADING:3 TRAILING:3 SLIDINGWINDOW:4:15 MINLEN:36; **(26) Trinity**: version 2.8.4, --seqType fq --full_cleanup --min_contig_length 250; **(27) GMAP:** version 2018-07-04, default parameters; **(28)** BLAT: version 36, default parameters; **(29) PASA:** --stringent_alignment_overlap 30.0 --MAX_INTRON_LENGTH 20000 –TRANSDECODER; **(30) EvidenceModeler**: --segmentSize 100000 --overlapSize 10000; **(31) TransposonPSI:** default parameters; **(32) InterProScan:** version v5.10–50.0, -goterms; **(33) PfamScan:** default parameters; **(34) tRNAscan-SE:** version 2.0, default parameters; **(35) RNAmmer**: version 1.2, -S euk -m lsu,ssu,tsu; **(36) snoscan:** version 0.9.1, default parameters; (**37) INFERNAL:** version 1.1.2, default parameters; **(38) BUSCO:** version 3.0.2, default parameters**; (39) LAI:** default parameters; (40) CD-Hit: version 4.8.1, -c 0.7.

## Acknowledgements

This work was supported by Yunnan Innovation Team Project and Natural Science Foundation of Yunnan (to L.-Z.G.).

## Author contributions

LZG conceived and designed the study; XGZ, CS and DZ contributed to the collection and preparation of the samples; WL and KL performed the genome assembly; WL performed genome annotation; QJZ performed data visualization; WL and LZG drafted the manuscript; LZG revised the manuscript.

